# High Resolution Epigenomic Atlas of Early Human Craniofacial Development

**DOI:** 10.1101/135368

**Authors:** Andrea Wilderman, Jeffrey Kron, Jennifer VanOudenhove, James P. Noonan, Justin Cotney

## Abstract

Defects in embryonic patterning resulting in craniofacial abnormalities are common birth defects affecting up to 1 in 500 live births worldwide, and are mostly non-syndromic. The regulatory programs that build and shape the craniofacial complex are thought to be controlled by information encoded in the genome between genes and within intronic sequences. Early stages of human craniofacial development have not been interrogated with modern functional genomics techniques, preventing systematic analysis of genetic associations with craniofacial-specific regulatory sequences. Here we describe a comprehensive resource of craniofacial epigenomic annotations and systematic, integrative analysis with a variety of human tissues and cell types. We identified thousands of novel craniofacial enhancers and provide easily accessible genome annotations for craniofacial researchers and clinicians. We demonstrate the utility of our data to find likely causal variants for craniofacial abnormalities and identify a large enhancer cluster that interacts with *HOXA* genes during craniofacial development.

## Introduction

Formation of the craniofacial complex is an intricate process of precisely timed events that occurs relatively early in vertebrate embryonic development. For example, in human embryonic development the majority of the events that lead to the formation of the human face and skull occur during the first ten weeks of gestation^1^. Defects in the orchestration of these events result in several different congenital abnormalities including failure of features to fuse (orofacial clefting) and premature fusion of structures (craniosynostosis). Worldwide, orofacial clefting is one of the most common birth defects, affecting ∼1 in 700 live births^2^. The majority of those affected with these types of clefting do not have defects in other tissues or organ systems and thus are referred to as “non-syndromic”^3^. While these birth defects are largely repairable through surgical means, the financial, sociological, and psychological effects have a much broader impact and represent a significant public health burden^4-7^. Screening, prevention, and non-surgical therapeutic options are thus highly desirable. The high heritability of such disorders suggests a major genetic component^8,9^; however, causative genetic changes have only been identified in a fraction of those affected^10^. Candidate gene approaches have identified mutations in seven different genes that explain less than ten percent of non-syndromic orofacial clefting cases^11^. In the past decade, several genome wide association studies, copy number variant analyses, and whole exome sequencing studies have sought to identify additional genetic sources of non-syndromic orofacial clefting^11-21^. These studies identified common and rare variants associated with orofacial clefting, but most are located in non-coding portions of the genome. Our genomes are littered with gene regulatory sequences, located primarily in intronic and intergenic sequences, that are active in a small number of tissues and/or developmental stages in humans^22^. While the regulatory potential of the human genome is still not completely understood, defects in regulatory sequences can cause non-syndromic developmental defects in humans and mice^23-26^. These findings, coupled with the non-syndromic nature of most orofacial clefting cases, suggest defective gene regulatory sequences may underlie much of the incidence of orofacial clefting. However, mapping of chromatin states and identification of craniofacial-specific regulatory sequences has been ignored by large functional genomics efforts such as ENCODE and Roadmap Epigenome^22^. The lack of craniofacial-specific gene regulatory information has impeded the identification of regulatory circuitry important for human craniofacial development and has prevented accurate interpretation of clinical genetic findings in patients with craniofacial disorders. Lastly, without sufficient biological context, prioritization and developing of hypotheses to test genetic associations with craniofacial abnormalities are hindered^27-30^. Here we present a comprehensive resource of functional genomics data and predicted chromatin states for important stages of early human craniofacial development. We have profiled multiple biochemical marks of chromatin activity in developing human craniofacial tissue samples encompassing 4.5 to 8 post conception weeks. We have comprehensively compared these data with publicly available genomic and genetic data from 127 epigenomes which include a wide variety of adult and fetal tissues. We provide annotations consistent with large consortia efforts^22^ in formats easily loadable into modern genome browsers to enable exploration by other researchers without large computational effort. We demonstrate how to mine this data for biological features relevant to craniofacial development and how to experimentally validate target gene interactions. In total, our analyses have identified thousands of previously unknown craniofacial enhancer sequences. These analyses will facilitate interpretation of genetic variation in the context of congenital craniofacial defects, and will enable future experimental testing of enhancer-target gene interactions in developing craniofacial tissues.

## Results

### Profiling of Histone Modifications in Developing Human Embryonic Craniofacial Tissue

Chromatin immunoprecipitation of post-translational histone modifications coupled with next generation sequencing (ChIP-Seq) is a powerful method to identify active regulatory sequences in a global fashion from a wide variety of biological contexts^22^. Many of the regulatory elements identified by this method are specific to the biological context queried^31,32^ (i.e. tissue type or developmental stage) and are enriched for genetic associations with disease in a relevant tissue (i.e. immune-related disorder associations in immune cell-specific enhancers)^33,34^. To identify regulatory sequences important for human craniofacial development, we utilized ChIP-Seq of six post-translational histone modifications across multiple stages and multiple biological replicates of early human craniofacial development. We focused our efforts on histone modifications both profiled by large consortia and strongly associated with multiple states of chromatin activity. We performed parallel ChIP-Seq experiments on craniofacial tissues obtained from 17 individual human embryos spanning the critical window for the formation of the human orofacial apparatus (Fig. 1a). Specifically, we profiled marks ranging from those associated with repression (H3K27me3), promoter activation (H3K4me3), active transcription (H3K36me3), and various states of enhancer activation (H3K4me1, H3K4me2, and H3K27ac) (Fig. 1b)^35^. We profiled at least three biological replicates for four distinct Carnegie stages (CS) (CS13, CS14, CS15, and CS17) encompassing 4.5 post conception weeks (pcw) to 6 pcw. We also profiled single biological samples from CS20 (8 pcw) and 10 pcw embryos (Fig. 1c). We obtained over 5.3 billion ChIP-Seq reads across a total of 106 datasets, with mean total reads and uniquely aligned reads per sample of 50.3 and 37.3 million respectively (**Supplementary Table 1**). Overall the samples correlated well by mark and stage of development (Fig. 2a **and Supplementary Fig. 1**). We uniformly processed these data to identify reproducibly enriched regions for each mark within each stage. The genomic features identified by each set of enriched regions closely mirror what has previously been reported for each of these post-translational marks (Fig. 2b and **Supplementary Fig. 2**)^32,35^. For example, we observed very strong enrichment of H3K4me3 at promoters of genes and identified a large number of intronic or intergenic regions enriched for H3K27ac. When we examined all the samples for a given Carnegie stage, we identified thousands of enriched regions, at each stage for each mark, that were found in at least two biological replicates (Fig. 2c).Combined, these results indicated our ChIP-Seq data from human embryonic tissues were of high quality, reflected the previously described nature of these marks, and was likely to identify tissue-specific regulatory sequences.

**Figure 1.**
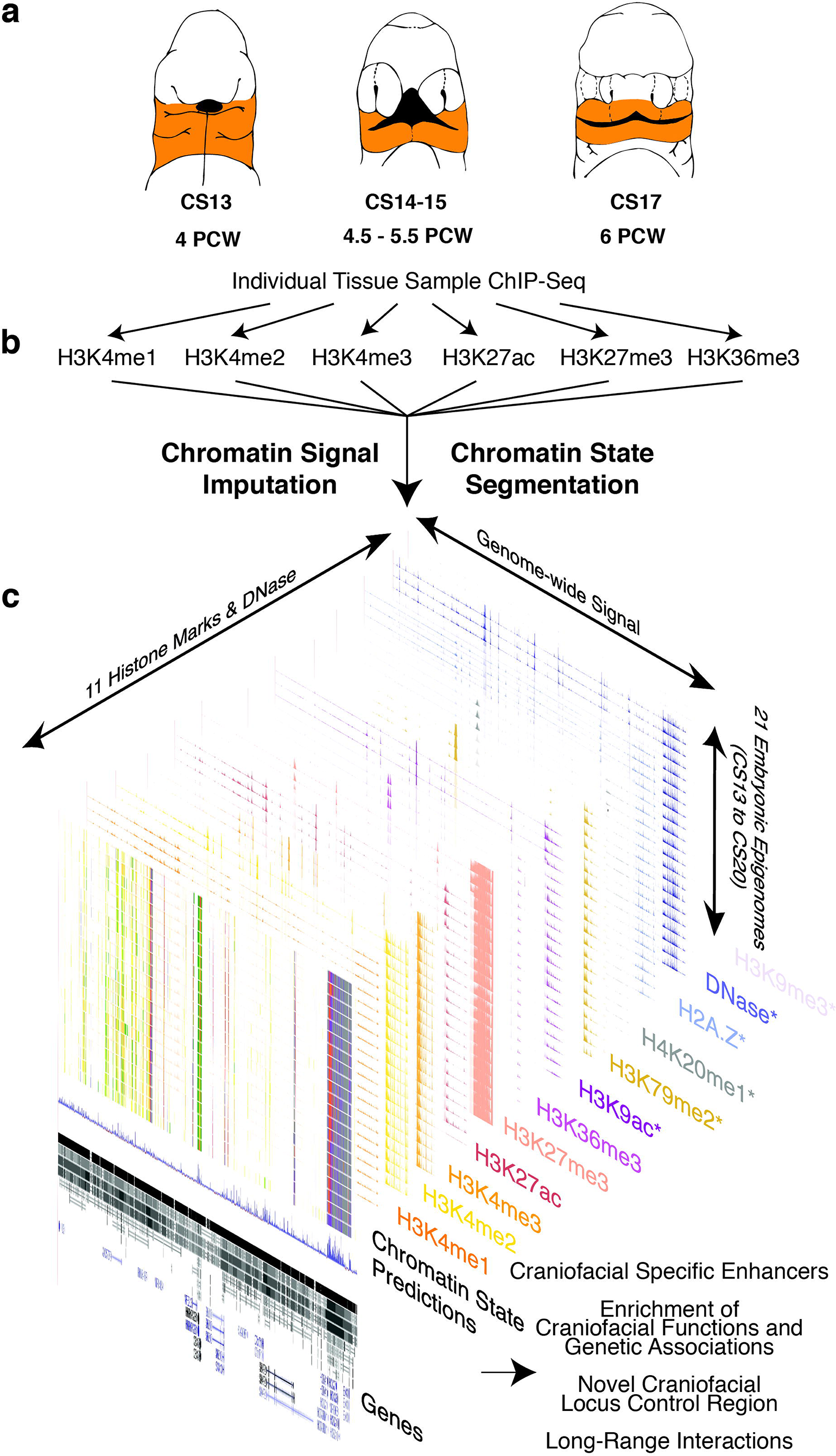
Overview of Epigenomic Profiling of Early Human Craniofacial Development. **a.** Stages and craniofacial tissues (orange shading) of human embryonic development sampled in this study indicated as Carnegie Stages (CS) or approximate post-conception weeks (pcw). Voids or cleavages in the embryo are indicated by black shaded regions. **b.** Six post-translational modifications of histones were profiled in parallel from individual human embryos via ChIP-Seq. **c.** Signals from primary ChIP-Seq data were imputed using ChromImpute^49^ to match the 12 epigenomic signals profiled by Roadmap Epigenome^22^. Asterisks indicate signals containing only imputed data. These imputed datasets were then used to predict chromatin states using a Hidden Markov Model approach (ChromHMM)^44^ across the genome for each craniofacial tissue sample. These chromatin states were then used for downstream functional analyses to determine relevance for craniofacial biology and disease.

**Figure 2.**
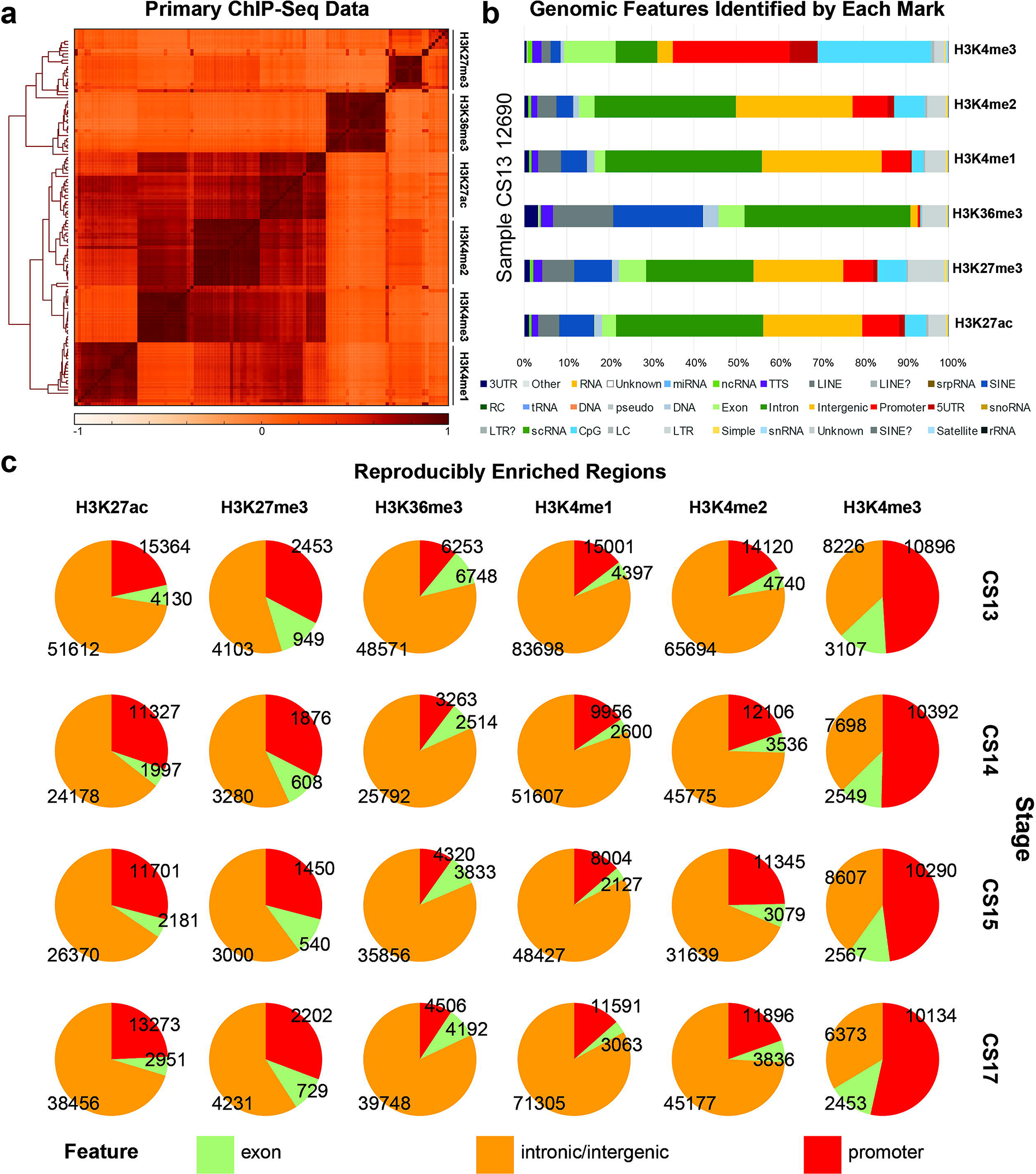
Histone Modification Profiles in Human Craniofacial Development. **a.** Heatmap and hierarchical clustering of pairwise Pearson correlations for non-overlapping 10kb bins across the human genome for 114 individual histone modification profiles from human craniofacial tissues. Relatedness of epigenomic profiles by sample indicated by dendrogram along vertical axes of heatmap. Darker orange indicates positive correlation between datasets. **b**. Genomic feature annotations identified by peak calls from six histone modification profiles from the same tissue sample plotted as cumulative percentage of total peaks. Peak enrichments and genomic annotations were performed using HOMER^83^. **c.** Histone modification peaks identified in at least two separate tissue samples from the same developmental stage and annotated into three broad categories: promoter (2kb upstream of TSS), exons, and all other intronic or intergenic locations.

### Generation of Human Craniofacial Chromatin State Segmentations

Defining enriched regions for a single histone modification such as H3K27ac has been utilized to identify active regulatory sequences from a variety of tissues, biological contexts, and different species^36-40^. However, in the absence of H3K27ac, other marks can identify active regulatory sequences, and low levels of H3K27ac may be present at enhancers that are either about to become active or are no longer active^41-43^. More advanced methods, such as using machine learning techniques and integrating multiple chromatin signals from a single tissue, allow segmentation of the genome into a more complex array of biological states^44,45^. These techniques can identify tissue-specific and disease-relevant regulatory information in a large cohort of tissues^35,46^. To leverage such available data to identify regulatory information likely to be critical for craniofacial development, we processed our data in a uniform fashion to match those generated by Roadmap Epigenome (Methods)^22^. Using p-value based signals^47,48^ for each of the six epigenomic marks we assayed, along with the same type of signals for 12 epigenomic marks for 127 tissues and cell types generated by Roadmap Epigenome, we imputed our data to create a uniform, directly comparable dataset^49^ (Fig. 1c). The imputed samples’ signals correlated well with their primary signals and clustered generally by mark and biological function (Fig. 3a and Supplementary Fig. 3). Using the imputed craniofacial data, we then segmented the genome for each embryonic sample based on previously generated models of 15, 18, and 25 states of chromatin activity^22^. We identified similar numbers and proportions of segments in each state in our tissues (Fig. 3b and Supplementary Fig. 4).The 25-state model results showed the most similar trends across these measures and utilized all of the primary data generated in our study when compared to those previously generated by Roadmap Epigenome (Fig. 3c,d and Supplementary Fig. 4); therefore we focused our downstream analyses on these segmentations. Using the 25-state segmentations, we reproducibly identified 75928 segments in at least one of six enhancer categories defined by Roadmap Epigenome (EnhA1, EnhA2, EnhAF, EnhW1, EnhW2, and EnhAc). To determine if these segmentations are enriched for craniofacial enhancers, we first turned to a large catalog of experimentally validated developmental enhancers tested in mouse embryos and available in the Vista Enhancer Browser^50^. We identified over 80% of all craniofacial-positive enhancers in this database. Moreover, our enhancer annotations were significantly enriched for craniofacial enhancers versus those that lacked craniofacial activity (p = 3.28 x 10^-14^) (Fig. 4a,b and Supplementary Fig. 5). While these results are encouraging - namely, that our data identified craniofacial enhancers - they did not reveal any specificity for craniofacial tissues in our chromatin state annotations. To address this problem, we quantitatively compared H3K27ac signals at all enhancer segments in our data with 127 samples from Roadmap Epigenome. Both hierarchical clustering and principal component analysis showed that our samples were well correlated with one another in this multi-tissue context (Fig. 4c and Supplementary Fig. 6). They were most similar to embryonic stem cells (ESC) and cell types derived from them (ESDR), but distinct from fetal and adult samples present in Roadmap Epigenome data. Previous analyses of Roadmap Epigenome have identified a significant number of enhancers that are tissue-specific^22^. To identify such novel enhancers in craniofacial tissue we first determined if any of our enhancer segments were ever annotated as such in the 127 samples obtained from Roadmap Epigenome. We identified 6651 enhancer segments (8.7% of total craniofacial enhancer segments) in our craniofacial epigenomic atlas that were never annotated as any type of enhancer state in all of Roadmap Epigenome (**Supplementary Table 2**). To determine if these sites are relevant for craniofacial development or represent spurious segmentations in our data we analyzed sequence content of these regions and functional enrichments of genes potentially regulated by these regions. When we assessed the novel craniofacial-specific enhancers for enrichment of transcription factor binding sites, we identified motifs matching those of *TWIST2*, *LMX1B*, *SIX1*, *NKX6.1*, multiple members of the *LHX* and *HOX* families, and *TCF12*, all of which have been implicated in craniofacial and skeletal development^51-57^ (Fig. 4d **and Supplementary Table 3**). Utilizing the Genomic Regions Enrichment of Annotations Tool (GREAT)^58^, we found significant enrichment of craniofacial-specific enhancers assigned to genes associated with craniofacial abnormalities such as cleft palate in both humans and mice (Fig. 4e **and Supplementary Fig. 7**). Interestingly, we also identified more general categories of enrichment amongst the putative gene targets including general transcriptional activators (**Supplementary Table 4**). When we interrogated this list of transcription factors, we found significant enrichment for expression in both craniofacial and appendicular skeleton (Fig. 4f). These results suggest that many of the novel craniofacial enhancers we identified are likely to play a direct role in patterning of the bones of the face, jaws, and portions of the skull. However, it is unclear whether they are directly involved in human craniofacial abnormalities.

**Figure 3.**
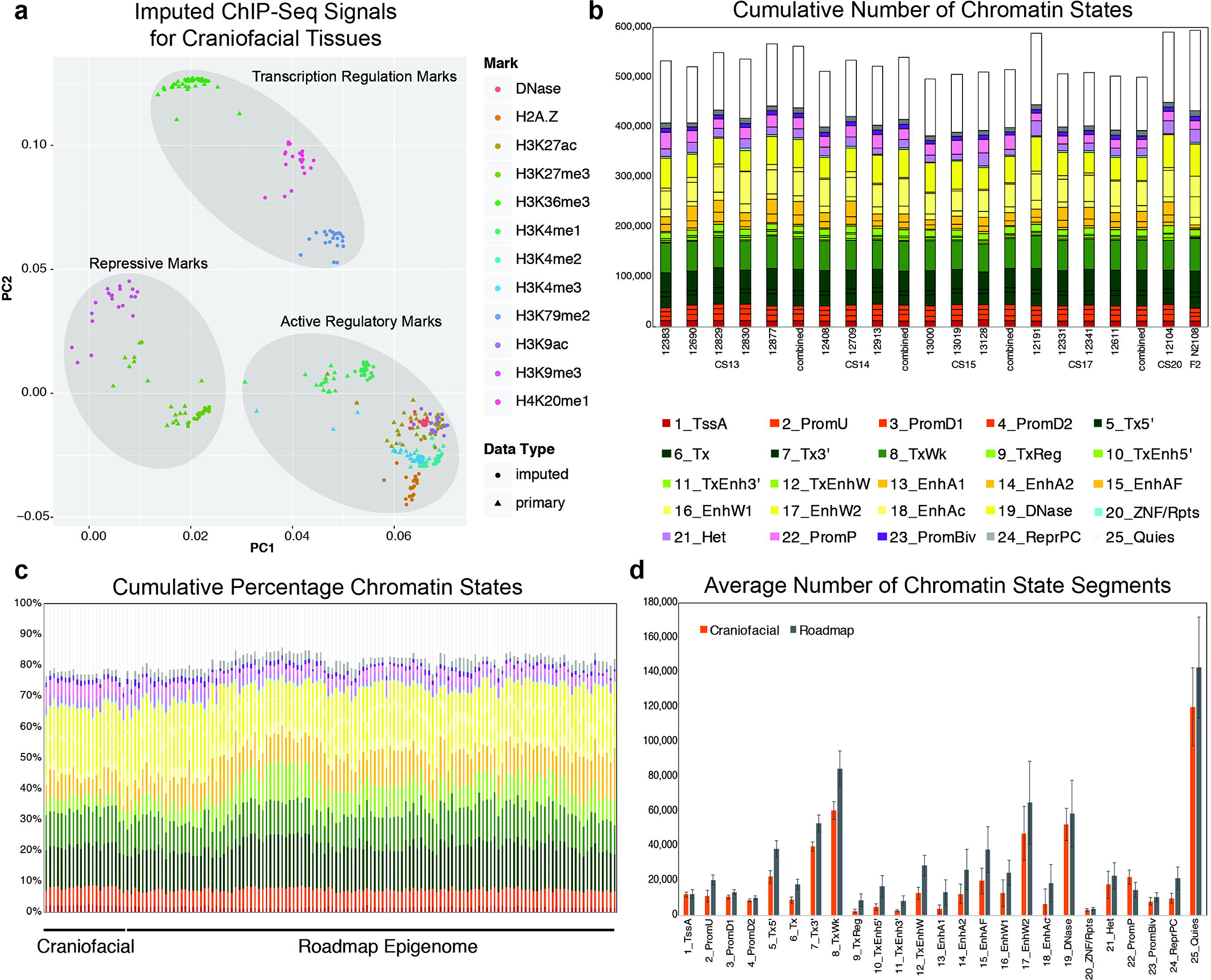
Imputation of Craniofacial Epigenomic Signals and Chromatin State Segmentation. **a.** Principal component analysis projection of first two component dimensions for 252 imputed and 114 primary epigenomic profiles for human craniofacial samples across non-overlapping 10kb bins. Samples are color coded by epigenomic mark and shapes indicate primary versus imputed data types. Samples generally cluster into three broad categories of activity: repression, regulatory element activation, and transcription regulation. **b**. Numbers of individual chromatin state segments identified by each of the color coded 25 states of chromatin activity based on imputed epigenomic signals for each of the 21 tissue samples profiled. **c**. Comparison of cumulative percentage of each chromatin state between craniofacial samples profiled here and 127 segmentations generated by Roadmap Epigenome^22^. **d**. Mean numbers of segments annotated in each of the 25 states across 21 craniofacial samples (orange) and 127 Roadmap Epigenomes (gray). Error bars represent standard deviation. Overall chromatin state segmentation in craniofacial samples identifies similar numbers and percentages of each of 25 states published by Roadmap Epigenome^22^.

**Figure 4.**
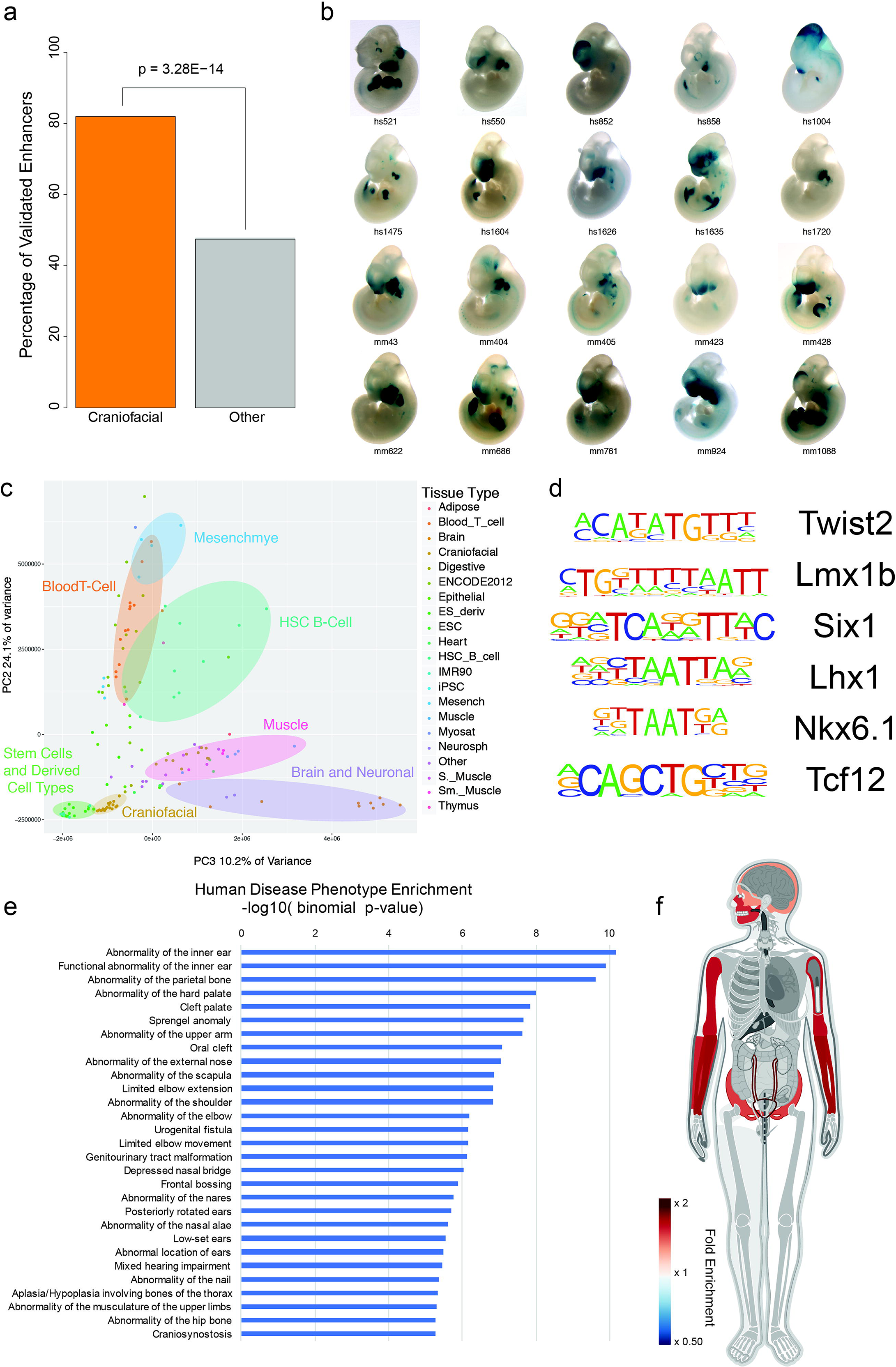
Chromatin State Segmentations Identify Novel Craniofacial Regulatory Sequences. **a.** Percentage of *in vivo* validated embryonic enhancers with (orange) or without (grey) craniofacial activity from the Vista Enhancer Browser^50^ identified by craniofacial chromatin segments annotated as enhancer states. Significance determined by Fisher’s exact test. **b.** Selected validated enhancers with craniofacial activity identified by this study from the the Vista Enhancer Browser. **c.** Principal component analysis projection of second and third component dimensions for 146 H3K27ac profiles at 425380 regions annotated as enhancer segments in any of the samples profiled here or Roadmap Epigenome. Samples are color coded by group annotations assigned by Roadmap Epigenome or craniofacial samples from this study. Percent of variance across samples explained by each component are indicated along each axis. **d.** Transcription factor position weight matrices identified by HOMER^83^ as enriched in novel craniofacial enhancer segments. **e.** Significant enrichments of human disease phenotypes for genes assigned to novel craniofacial enhancer segments as reported by GREAT^58^. **f.** Enrichment of anatomical expression of transcription factors identified as potentially regulated by novel craniofacial enhancer segments as reported by GeneORGANizer^88^. Heat indicates fold enrichment of expression in individual anatomical region or organ. Craniofacial and appendicular skeleton showed most significant enrichments.

**Figure 5.**
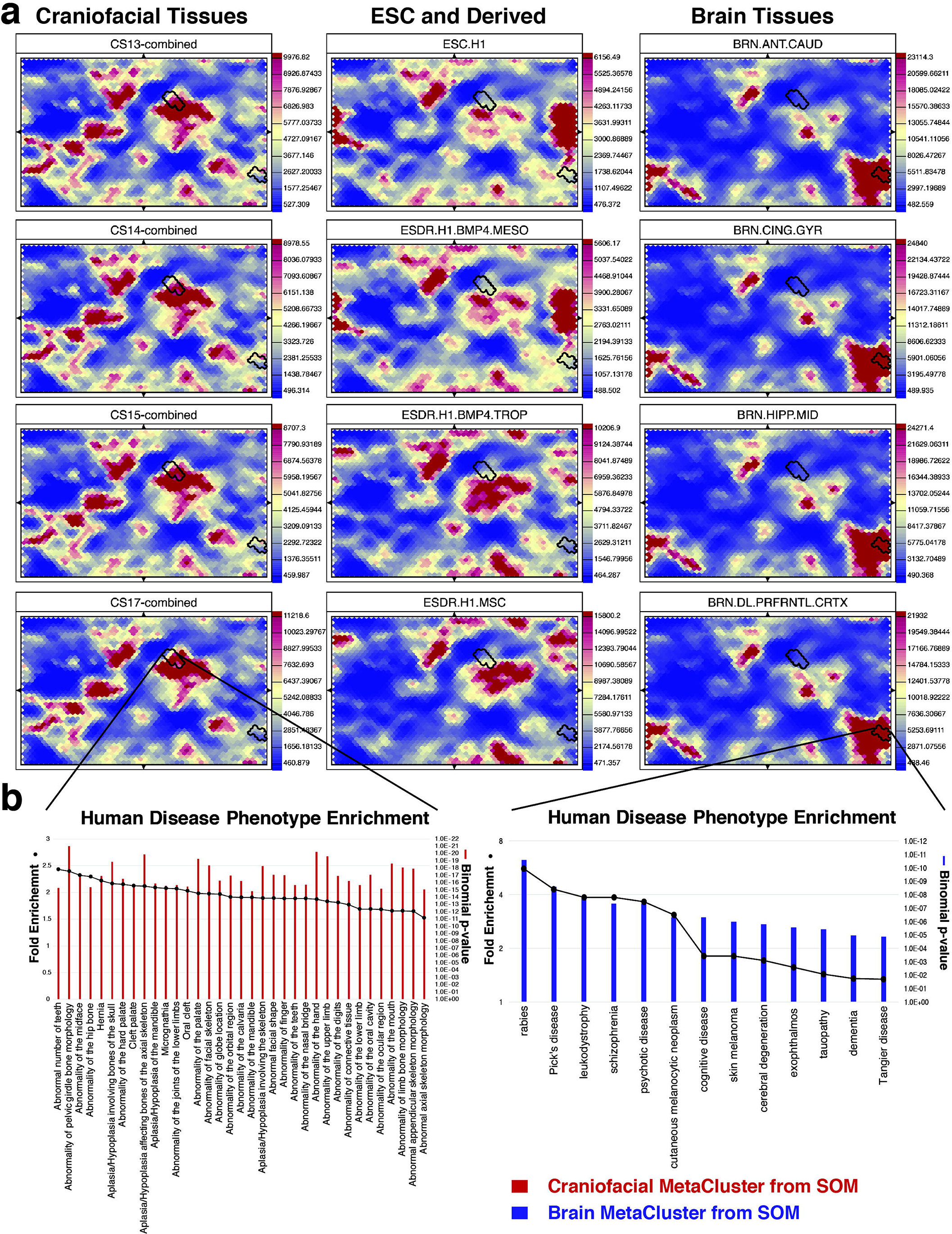
Self-Organizing Map for Biological Mining of Craniofacial Enhancers. **a.** Flattened projections of toroid self-organizing map generated from H3K27ac signals from 146 samples across 425380 enhancer segments consisting of 2500 individual hexagonal units for four craniofacial tissues, four embryonic stem-cell and related cell types, and four adult brain tissues. Higher scoring units in a given tissue are indicated by red, lower scoring units by blue. Two selected metaclusters scoring highly for craniofacial or brain tissues are indicated by black outlines. **b**. Fold enrichment (dots) and significance (bars) of top human disease phenotypes associated with genes assigned to enhancer segments by GREAT^58^ in each metacluster. A metacluster highly scoring reproducibly in craniofacial tissues is enriched for enhancers putatively assigned to genes associated with a wide variety of craniofacial abnormalities. A metacluster highly scoring across brain tissues is enriched for diverse brain and neurological diseases. While PCA and hierarchical clustering identified craniofacial tissues were more similar to ESC and ESC-derived cell types, the self-organizing map identifies distinct clusters of enhancers specific to craniofacial tissues.

**Figure 6.**
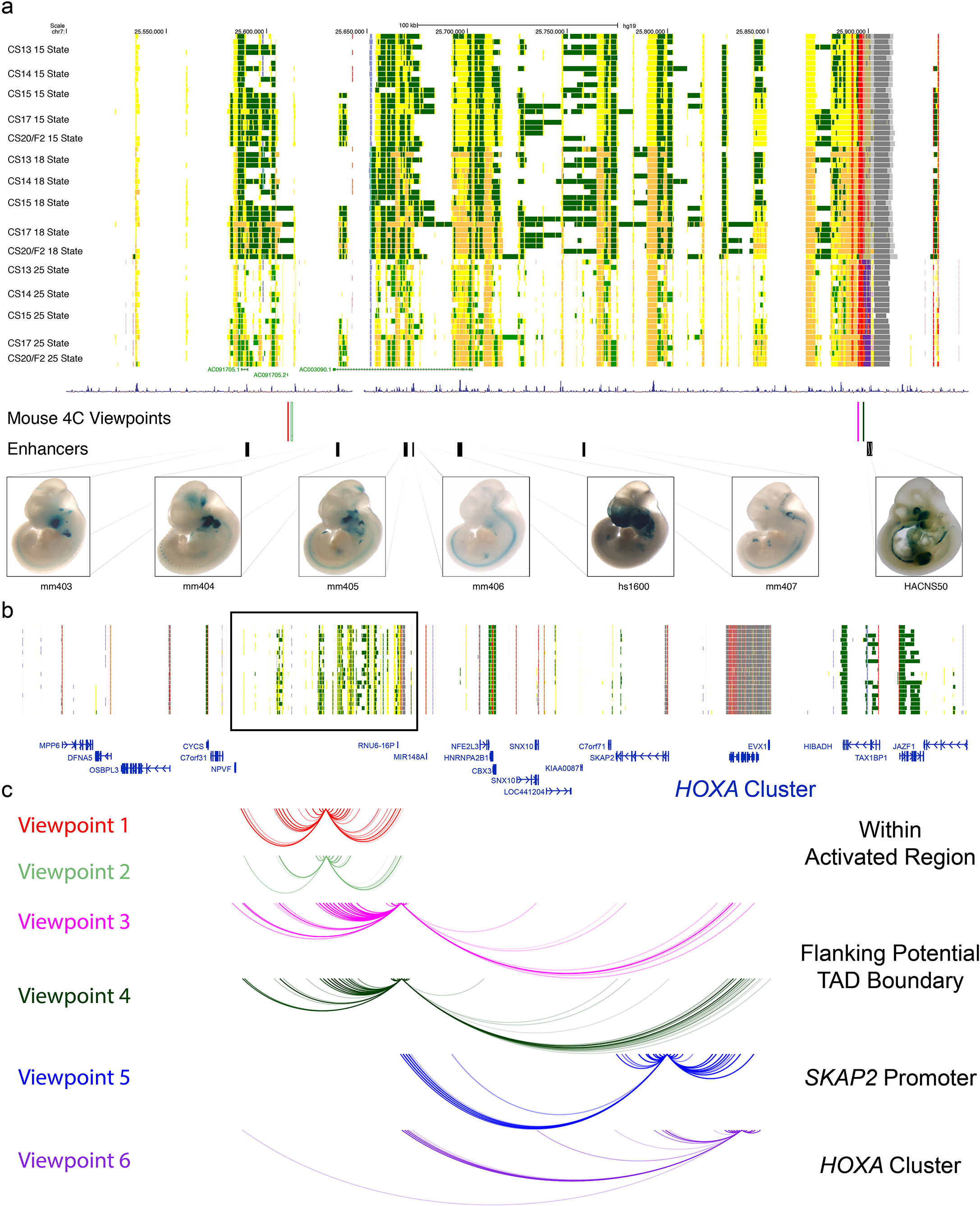
Identification of Potential Craniofacial Locus Control Region for *HOXA* Gene Cluster. **a.** Large 450kb window lacking any annotated protein-coding genes with extensive enrichment of activated enhancer (yellow and orange) and transcriptionally active (green) segment annotations in human craniofacial tissue. See Figure 3b for full annotations. Multiple validated craniofacial enhancers have been identified in this window by the Vista Enhancer Browser. In this study we tested and validated the craniofacial enhancer activity of HACNS50, located within the bivalent chromatin state at the right of the displayed window. Segments interrogated by 4C-Seq indicated by vertical colored viewpoint bars **b.** Approximately 3Mb window of the human genome encompassing the window identified in panel **a** (black box) and containing the *HOXA* gene cluster. **c.** Spidergrams indicating significant interactions between color-coded viewpoints and distal sites identified by 4C-Seq in mouse E11.5 craniofacial tissue. Viewpoints 1 and 2 do not cross putative TAD boundary near HACNS50 enhancer. Viewpoints 3 and 4 make significant contacts within identified window and with the *HOXA* gene cluster. Reciprocal experiments from the *HOXA* gene cluster (viewpoint 6) indicated significant long-range interactions with both boundaries of the window in panel **a**.

To begin to explore this uncertainty, we turned to genome wide association data obtained from the GWAS catalog related to orofacial clefting and craniofacial morphology^17,21,59-63^. We overlaid associations from these studies with each of the segmentation maps from our data, as well as data from Roadmap Epigenome, and assessed enrichment. We observed significant enrichment of orofacial clefting tag SNPs in most of our craniofacial samples and relatively few Roadmap Epigenomes (**Supplementary Fig. 8a**). These analyses identified several enhancer segments that directly contain strong genetic associations. For instance, we identified a discrete enhancer state in the noncoding region between *IRF6* and *DIEXF* that contains a tag SNP previously associated with non-syndromic cleft lip and palate^64^ (**Supplementary Fig. 8b**). This particular region can directly influence *IRF6* expression and is potentially a causative allele for orofacial clefting^65^.We also identified 13 other regions that are identified in craniofacial tissue and directly contain such tag SNPs, including an intronic sequence of the *TXNDC16* gene^59^ (**Supplementary Fig. 8c and Supplementary Table 5**). These findings suggest that our chromatin state maps will be extremely useful in identifying and prioritizing causative variation in patients affected by craniofacial abnormalities.

### Machine learning approaches to mining of activated craniofacial enhancer data

To more comprehensively explore our data for regions likely to be important for craniofacial development and human disease, we turned to the unsupervised machine learning method known as self-organizing maps. This approach is a powerful means to identify relationships within large genomic datasets, but also allows fine-grained analysis relevant to specific biological questions^66^. We first extracted H3K27ac signals from all of our craniofacial samples and all Roadmap Epigenome samples across all enhancer segmentations, resulting in signal measurements for 425000 enhancer segments in 146 epigenomes.The resulting matrix was used to train a self-organizing toroid map with a map size of 2500 units; we selected the best scoring map from 50 map building trials. We then clustered each of the units of the map into metaclusters and found 199 that identify enhancer segments that have similar signal properties and are likely to be biologically related (Fig. 5a). For each enhancer segment, we assigned potential target genes and overlaid the gene assignments to each unit. Based on these gene assignments, we then determined the gene and human phenotype ontology enrichments of each unit. This resource is available for interrogation via a standard web browser, allowing for retrieval of regions, genes, and functional associations for each unit and metacluster. Inspection of this map identified several metaclusters that showed distinct H3K27ac activation in craniofacial tissues. These clusters were enriched for a number of ontologies related to craniofacial biology and abnormalities. For example, we identified a metacluster that showed significantly increased H3K27ac signal in craniofacial samples relative to other tissue types and that is enriched for potential target genes associated with various craniofacial abnormalities (Fig. 5b). We obtained similar types of functional enrichments when performing k-means clustering directly on the matrix of H3K27ac signals using the same number of clusters utilized for the self-organizing map (**Supplementary Fig. 9a and Supplementary Table 6**). When we assessed the sequence content of clusters most specific for craniofacial activity, we identified enrichment of motifs for the *ALX*, *DLX*, *HOX*, and *MSX* families of transcription factors (**Supplementary Fig. 9b**).

### Identification of novel craniofacial locus control region and potential regulatory targets

Thus far, our analyses have focused on the annotation and activation state of individual genome segments in bulk. However, these enhancers likely do not operate in isolation and clusters of enhancers activated in concert have been shown to be powerful regulators of important genes for a given tissue or cell type^67^. To identify such enhancer clusters, we applied a sliding window approach to detect enrichment of craniofacial enhancer states relative to both randomly chosen sequences as well as those identified by Roadmap Epigenome. We identified 582 regions across the genome that demonstrate high levels of craniofacial enhancer activity (**Supplementary Table 7**). These windows had an average size of ∼400kb but ranged up to 2 Mb in length. In many cell types these clusters of enhancers, sometimes referred to as super enhancers, are embedded in the genome both surrounding and within the introns of their likely tissue-specific target^68^. Indeed, most of the windows we identified contained multiple genes and were enriched for developmental genes, including multiple *Frizzled*, *WNT*, *ALX*, *DLX*, and *TBX* family members (**Supplementary Table 8**). Interestingly, we identified 37 large windows that were located entirely in intergenic space and did not overlap a promoter region for any known genes. These windows represent potentially novel large clusters of regulatory regions, but their targets and activities are difficult to interpret using linear genomic annotations and distances. Given that studies have shown that our genome can form numerous long range interactions^69,70^, especially between regulatory regions, we sought to determine if any of these intergenic clusters of putative enhancers could be important for craniofacial development by identifying direct three-dimensional interactions in relevant tissues. To ensure that we were interrogating bona fide enhancer clusters, we focused our downstream efforts on regions that contained *in vivo*-validated craniofacial enhancers. We identified a single window encompassing a 450kb region located on chromosome 7 that contains five confirmed craniofacial enhancers from the Vista Enhancer Browser^50^ (Fig. 6a). This region also contains a unique chromatin signature at its 3’ end, where strongly active and repressed states are directly adjacent. This is most commonly observed at looping or topological domain boundaries. Indeed, long-range contact maps from human umbilical vein endothelial cells^69^ indicate this observed chromatin state transition is a topologically associated domain (TAD) boundary (**Supplementary Fig. 10**). We tested an element annotated as a bivalent chromatin state, which is highly conserved across mammals, near this chromatin boundary for enhancer activity^71^. It displayed strong craniofacial and limb enhancer activity in the E11.5 mouse embryo (Fig. 6a). Inspection of chromatin data from Mouse ENCODE^72^ indicate similar patterns of activation in the orthologous window, suggesting functional conservation of chromatin state in this large region (**Supplementary Fig. 11**). Having demonstrated that this region is enriched for craniofacial enhancers and active chromatin states, we sought to determine the gene(s) this region interacts with and potentially regulates. This region is located between the *NPVF* and *NFE2L3* genes, neither of which appears to be active based on observed chromatin states in developing human craniofacial tissue. The next closest target is *CBX3*, which is strongly expressed in most cell types and has similar chromatin states in both our tissues and in Roadmap Epigenome. Comparisons of the mouse and human genomes revealed this window is part of a large syntenic block between the two species which stretches nearly 10 Mb in length with the *HOXA* gene cluster at its center (**Supplementary Fig. 12**). The enrichment for craniofacial enhancer annotations, harboring of six *in vivo* validated craniofacial enhancers, potential TAD boundary, and conservation both at the sequence and epigenomic level suggest this region is an important regulatory hub.

Two control regions, the early limb control region (ELCR) and the global control region (GCR), have been identified for the *HOXD* gene cluster that are important for regulation of the cluster’s expression in the developing mammalian limb. The exact coordinates of the ELCR are unknown, but they are thought to be located in the large noncoding region adjacent to the cluster, while the GCR is approximately 250kb away, beyond the *LNP* gene^73,74^. No such control regions have been identified or described for the *HOXA* cluster. Furthermore, loss of at least one gene in the cluster, *HOXA2,* has been implicated in cranial neural crest skeletal morphogenesis and results in mice born with cleft palates and other craniofacial abnormalities^57,75^. The region we have identified is located nearly 1.5 Mb from the *HOXA* cluster and contains at least seven annotated genes in the intervening genomic sequence; thus it is not clear whether this region could regulate the *HOXA* gene cluster. Utilizing circularized chromosome conformation capture with sequencing^76^ (4C-seq) we assessed the interactions of four viewpoints in this window in E11.5 mouse craniofacial tissue. For two viewpoints, we identified extensive interactions within the identified window that do not cross the putative TAD boundary. When we assessed viewpoints flanking the TAD boundary, one of which contained the active enhancer HACNS50^71^, we observed interactions within this identified region as well as significant interactions with the *HOXA* gene cluster (Fig. 6b). To confirm these interactions, we performed additional 4C-seq experiments utilizing viewpoints located directly within the *HOXA* cluster and the promoter of the *SKAP2* gene. We observed strong interactions between both of these viewpoints and the TAD boundary of the original window. Interestingly, *HOXA* made contacts with the outer limits of this window but not within the window. These findings illustrate that the region we identified in human craniofacial tissue makes strong contacts over nearly 1.5 Mb with genes of the *HOXA* cluster in developing mouse craniofacial tissue and indicate it could be a conserved global control region important for craniofacial development.

## Discussion

Our understanding of the regulation of craniofacial development and the genetic changes that give rise to developmental defects has not advanced greatly in the last decade despite it being a heavily studied area of human and mouse biology and the advent of more advanced genomic technologies. Recent large consortia efforts to identify the genetics of common disease have gained traction utilizing tissue-specific annotations of the genome to identify potential regulatory regions and overlaying genetic associations^33,34^. Such genetic association data exist for craniofacial abnormalities, but the lack of craniofacial-specific annotations of regulatory function have prevented systematic identification of causal genetic changes. We have addressed this need by generating an extensive resource of functional genomics data obtained directly from human craniofacial tissues during important stages of formation of the orofacial apparatus. We have uniformly processed our data to allow integration of these data with similarly generated signals from a variety of human tissues and developmental stages. These analyses have allowed us to generate craniofacial-specific annotations of chromatin states across the human genome. These chromatin state segmentations reveal tens of thousands of regions with potential gene regulatory activity in craniofacial development. Over 6000 of the enhancer segments we identified have never been annotated previously as having enhancer activity in 127 different cell types. These regions are strongly enriched near genes implicated in craniofacial development and would have remained unknown to craniofacial researchers relying solely on the current state of genome annotations. Indeed, recent targeted sequencing of GWAS intervals at 13 loci in patients affected by craniofacial abnormalities likely excluded important craniofacial regulatory regions due to the lack of appropriate chromatin state annotations^77^ (**Supplementary Fig. 13**). These findings illuminate that our current understanding of the regulatory information our genomes encode is incomplete and reinforces the need for more and higher resolution tissue-specific chromatin state annotations.

To illustrate the utilization of this resource we analyzed these data in many different fashions to narrow down particular regions of interest and to interrogate them for genetic and functional associations with craniofacial development. Furthermore, we demonstrated that once a region of interest has been identified, it is possible to develop a hypothesis of potential gene regulatory targets and directly test them *in vivo* in the context of both genomic and functional conservation in the mouse. Here, we chose to focus on clusters of craniofacial enhancer segments that have been functionally verified in the developing mouse embryo to ensure relevance for craniofacial biology. Our windowing approach identified an extremely dense, large array of craniofacial enhancers, suggesting we have identified an important regulatory hub. The conservation of activating histone modification signals in developing mouse craniofacial tissues indicates this region is likely important for the formation of craniofacial features in multiple species. Additionally, the identification of direct long-range interactions between portions of this unique enhancer region, including a rapidly evolving conserved non-coding sequence (HACNS50)^71^, with the *HOXA* gene cluster suggest this region could be important not only for normal craniofacial development but also for evolution of the human skull. Lastly, this region has been implicated as a uniquely deleted segment in a patient with facial dysgenesis^78^ (**Supplementary Figure 10**). This patient was noted to have overtly normal organs and brain activity despite lacking most features of a face resembling other non-syndromic abnormalities caused by regulatory sequence defects^23-26^. Further genetic dissection of this region in cultured human cells or in the developing mouse are needed to determine the role this region plays in regulating this conserved cluster of *HOXA* genes.

We provide all our craniofacial functional genomics data and resulting chromatin state segmentations in several standard formats as well as a complete catalog of tracks that can be easily loaded into many modern genome browsers. Additionally, we provide our self organizing map of active enhancers across 146 samples as a website that can be explored by a variety of researchers to interrogate the regions, genes, and phenotypes relevant for their research without high level computational processing or expertise. (https://cotneylab.cam.uchc.edu/∼jcotney/CRANIOFACIAL_HUB/Craniofacial_H3K27ac_SOM/). This will allow the craniofacial community to develop hypotheses related to craniofacial abnormalities which are rooted in craniofacial biology instead of using chromatin state annotations from other tissues not directly related to the tissue of interest. These resources stand to bring the craniofacial research world firmly into the functional genomics era, advance our understanding of these disorders, and provide tools for clinicians seeking to diagnose patients utilizing whole genome sequencing.

## Acknowledgements

We would like to thank the donors to HDBR, as without them this resource would not be possible. We would also like to thank Camden Jansen and Ali Mortazavi for assistance in generation of the self-organizing map resource and Heather Adinolfi for technical assistance in the transgenic enhancer assay. Funding support was provided by an R00 award from NIDCR to J.C.

### Author Contributions

All ChIP experiments were performed by J.C. Sequencing libraries and sequencing runs were performed by J.K. Data analysis was performed by J.C. 4C-Seq experiments and analysis were performed by A.W. Writing and data interpretation were performed by all authors.

### Data and Code Availability

All data can be visualized in the UCSC Genome Browser using track hub functionality. Hub files and interesting browser examples can be found on our website: http://cotney.research.uchc.edu/data/

ChIP-Seq signals, peak calls, chromatin state segmentations and 4C-Seq data are available at GEO accessions GSE98251 and GSE97752.

All generic scripts used in processing ChIP-Seq and generating chromatin states are available on github: https://github.com/cotneylab/ChIP-Seq

All generic scripts for processing of 4C-Seq data from mouse are available on github: https://github.com/cotneylab/Mouse-HOXA-4C-Seq

## Methods

### Tissue Collection and fixation

Use of human fetal tissue was reviewed and approved by the Human Subjects Protection Program at UConn Health. Human embryonic craniofacial tissue was collected, staged and provided by the Joint MRC/Wellcome Trust Human Developmental Biology Resource (www.hdbr.org). Tissues were flash frozen upon collection and stored at -80⁏. Fixation for ChIP-Seq was performed as described in Cotney and Noonan, 2015^79^. Briefly, each tissue sample was rapidly thawed in 1 mL of ice cold phosphate buffered saline (PBS) and briefly homogenized with a disposable plastic pestle in a 1.5 mL microcentrifuge tube. Samples were then fixed by the addition of formaldehyde to a final concentration of 1% and incubated at room temperature on a rotisserie for 15 minutes. Samples were then quenched with 150 mM glycine at 10 minutes at room temperature. Tissue was collected by centrifugation (5 min, 2500g, 4⁏) and washed with 1 mL of fresh PBS. Fixed tissue pellets were then rapidly frozen in a dry ice/alcohol bath and stored at -80⁏ until batch processing for chromatin immunoprecipitation (ChIP).

### Antibody Specifications

Antibodies used in this study: anti-H3K27ac (ab4729, Abcam), anti-H3K4me1 (ab8895, Abcam), anti-H3K4me2 (ab7777, Abcam), anti-H3K4me3 (ab8580, Abcam), anti-H3K27me3 (07-449, EMD Millipore), anti-H3K36me3 (ab9050, Abcam).

### ChIP-Seq

Fixed tissue pellets were processed for ChIP as previously described^79^. Briefly, samples were thawed in 1 mL of 1x Cell Lysis buffer and incubated on ice for 20 minutes. Cells were lysed with dounce homogenization and nuclei were collected by centrifugation (5 min, 2500g, 4⁏). Nuclei were resuspended in 300 µL of 1x Nuclear Lysis buffer + 0.3% SDS + 2 mM sodium butyrate and incubated on ice for 20 minutes. Chromatin was sheared with a Qsonica Q800R1 sonicator system operating at amplitude 20 and 2⁏ for 30 minutes (10 seconds duty, 10 seconds rest). Samples were cleared by centrifugation (5 min, 20,000g, 4⁏) and soluble chromatin was transferred equally into six separate tubes with 10% reserved as an input control. SDS concentration was reduced to 0.18% with ChIP Dilution buffer. Protein G Dynabeads (ThermoFisher) separately preloaded with 2 µg of antibodies listed above were added to each chromatin aliquot. ChIP samples were incubated overnight at 4⁏ on a rotisserie. Chromatin was then immunoprecipitated on a magnet and supernatant was discarded. Beads were washed 8 times with 1 mL of 500 mM LiCl ChIP-Seq Wash Buffer and once with 1 mL of TE. Chromatin was eluted from the beads twice with ChIP Elution buffer at 65L for 10 minutes with constant agitation. Combined eluates for each ChIP were subjected to crosslink reversal overnight at 65⁏. Samples were then sequentially treated with RNAse A and proteinase K, purified with a PCR Purification Kit (Qiagen), and eluted in 50 uL of EB. ChIP samples were then quantified with picoGreen (ThermoFisher) and prepared for sequencing on Illumina instruments using the Thruplex 48S Library Prep kit (Rubicon Genomics) according to manufacturer’s instructions. Final libraries were quantified by QPCR (NEBNext Library Quant Kit for Illumina), multiplexed, and sequenced for 75 cycles across multiple flow cells on an Illumina NextSeq 500 instrument.

### Primary ChIP-Seq Data Analysis

Sequencing data was directly retrieved from Illumina’s Basespace Cloud service using Basemount command line tools provided by Illumina. Multiple FASTQs for each ChIP were combined and assessed for quality using FASTQC (v0.11.2)^80^ and compared visually using MultiQC (v0.9)^81^. Reads were then aligned to the human genome (hg19) using Bowtie2 (v2.2.5)^82^ keeping only uniquely mapped reads. Fragment sizes of each library were estimated using PhantomPeakQualTools (v.1.14)^47^. Histone modification enriched regions were identified and annotated using HOMER (v4.8.3)^83^. Reproducibly enriched regions were determined by creating a union of all enriched regions for a respective histone modification from all replicates of a single Carnegie stage and filtering for regions identified in at least two biological replicates using BEDtools (v2.25.0)^84^. We then generated p-value based signal tracks relative to appropriate input controls based on estimated library fragment size using MACS2 (2.1.1.20160309)^48^. All signal and enriched region files were converted for display in the UCSC Genome Browser using the Kent Source Tools (v329)^85^. Correlations of ChIP-Seq signals and Principal Component Analysis across samples and marks were calculated in non-overlapping 10kb windows using deepTools2 (v2.5.0.1)^86^.

### Roadmap Epigenome Data Retrieval

Aligned and consolidated primary ChIP-Seq reads in tagAlign format were retrieved from Roadmap Epigenome for eleven epigenomic signals: H2A.Z, H3K4me1, H3K4me2, H3K4me3, H3K9ac, H3K9me3, H3K27ac, H3K27me3, H3K36me3, H3K79me2, and H4K20me1. (http://egg2.wustl.edu/roadmap/data/byFileType/alignments/consolidated/). To ensure the most compatible signals with our data, p-value signals were generated by MACS2 from these data based on library fragment sizes reported by Roadmap Epigenome as above. DNase p-value signals were retrieved directly from Roadmap Epigenome (http://egg2.wustl.edu/roadmap/data/byFileType/signal/consolidated/macs2signal/pval/) and converted from bigWig to bedGraph for use with ChromImpute^49^ using Kent Source Tools ^85^. Chromatin state segmentations for 127 epigenomes and associated 15-, 18-, and 25-state model files were retrieved from Roadmap Epigenome (http://egg2.wustl.edu/roadmap/data/byFileType/chromhmmSegmentations/ChmmModels/).

### Chromatin Imputation

Bedgraph files for all p-value signals from primary ChIP-Seq data were converted to 25 bp resolution and processed for model training and generation of imputed signals for all samples using ChromImpute (v1.0.1) as previously described^49^. Resulting imputed signal tracks were converted to bigWig format for display in UCSC genome browser and converted to combined signal format at 200 bp resolution for use with ChromHMM (v1.12)^44^ using deepTools2^86^.

### Chromatin State Segmentation

Signal files for individual chromosomes for each craniofacial epigenome were binarized and segmentation was performed using previously published joint 15-, 18-, and 25-state chromatin models using ChromHMM as previously described^22^. Following segmentation, annotation of states and generation of genome browser files was performed based on annotations provided by Roadmap Epigenome. Individual models of 15, 18 and 25 chromatin states were also learned for each craniofacial epigenome using default settings in ChromHMM. Pearson Correlations and Principal Component Analyses were performed on total H3K27ac signals extracted observed in all imputed p-value signal tracks for craniofacial and Roadmap Epigenome samples from the union of all enhancer state segmentations (EnhA1, EnhA2, EnhAF, EnhW1, EnhW2, and EnhAc) using deepTools2^86^. All plots were made using tabular data generated by deepTools2 in R (v3.3.3)^87^.

### Functional Enrichments in Craniofacial Epigenomes

Craniofacial enhancer state segmentations (EnhA1, EnhA2, EnhAF, EnhW1, EnhW2, and EnhAc) were interrogated for tissue activity in the developing mouse embryo from the Vista Enhancer Browser^50^. Significance of overlap of enhancers identified in human craniofacial tissue and shown to be active in mouse craniofacial tissue relative to all other tissue annotations was determined using Fisher’s exact test. To identify totally novel craniofacial enhancers, enhancer state segmentations for all craniofacial segmentations were interrogated for single base overlap with the same states from all Roadmap Epigenomes using BEDtools^84^. These novel craniofacial enhancer segmentations were assessed for gene ontology and functional enrichments based on assigned target genes using GREAT (v3.0.0)^58^. Genes identified as transcriptional regulators by GREAT were assessed for enrichment of anatomical expression using default parameters in GeneORGANizer^88^. Sequence from novel craniofacial enhancer segmentations was extracted from hg19 using fastaFromBed within BEDTools^84^. The resulting sequences were assessed for transcription factor motif enrichment using HOMER^83^. Enhancer state segmentations from craniofacial epigenomes and all Roadmap epigenomes were interrogated for significance of overlap with GWAS tag SNPs associated with orofacial clefting and craniofacial morphology^17,21,59-63^ obtained from the GWAS Catalog (retrieved 2017-02-20)^89^ using Fisher’s exact test within BEDTools^84^.

### Self-Organizing Maps of Enhancer Activation

The self-organizing map of H3K27ac signal at all enhancer segments was generated as previously described^66^. Briefly, a union of all enhancer segmentations from craniofacial tissues and all samples in Roadmap Epigenome was generated and merged to form a consistent annotation of enhancers across the entire genome resulting in 425380 individual enhancer segments. H3K27ac signals from imputed p-value signal tracks for each of the 146 epigenomes were extracted for each of the 425380 enhancer segments. This matrix was then used to train a self-organizing map with 50 rows and 50 columns (2500 units) to allow for the possibility of small numbers of highly tissue-specific enhancers (<200) to be clustered together. We performed 50 training trials and retained the best scoring map. For this final self-organizing map we then annotated each unit with Ensembl (v75) genes based on association rules defined by GREAT^58^. Based on these unit/gene assignments we then determined enrichment of gene ontologies (http://geneontology.org/ontology/go.obo) and human phenotype ontologies from the Monarch Initiative^90^ (http://purl.obolibrary.org/obo/hp.obo) as previously described^91^. Clusters of units, or metaclusters, were then determined with four separate trials testing for the presence of up to 250 metaclusters as previously described^66^. The algorithm converged on 199 clusters as optimal for the self-organizing map generated above. Metaclusters were then assessed for functional enrichments as was done for individual units above. Metaclusters identified as specific for craniofacial and brain tissues were visualized using a JavaScript web-based viewer of the self-organizing map available here: https://cotneylab.cam.uchc.edu/∼jcotney/CRANIOFACIAL_HUB/Craniofacial_H3K27ac_SOM/

### K-means clustering of Enhancer Activation

K-means clustering of the same H3K27ac signal matrix utilized for the self-organizing map was performed using Cluster (v3.0)^92^. Rows were centered on the mean value of the row and normalized, the number of metaclusters identified in the self-organizing map analysis above was used as the k parameter, and 100 runs were performed. The clustering result was then visualized and craniofacial-specific clusters were extracted using Java TreeView^93^. Sequences underlying the enhancers in the craniofacial-specific clusters were extracted as above for novel craniofacial enhancers. We performed motif enrichment within these sequences using a combination of multiple tools for more robust enrichment determination^94^. Functional enrichment for these enhancers was determined as above using GREAT^58^.

### Identification of Enhancer Clusters

To identify clusters of craniofacial enhancers we first generated overlapping 200kb windows with a 50kb step size^84^. Next, we intersected these windows with all enhancer chromatin state segmentations from craniofacial tissues. We then calculated the fraction of each window annotated as an enhancer state. We tested for enrichment of enhancers in each window using permutation testing by randomly shuffling the craniofacial enhancer segments across the genome 1000 times using BEDtools^84^ and determining the fraction of each window annotated as an enhancer. Overlapping windows of significant enrichment were merged into a single contiguous region. Final enriched regions were assessed for overlap with gene annotations and validated craniofacial enhancers using BEDtools^84^.

### Transgenic Enhancer Assay

A 2.6 kb segment centered on the conserved sequence corresponding to HACNS50^71^ was amplified from human genomic DNA by polymerase chain reaction (PCR) using the following primers: HACNS50 F 5’-CACCCCATTTCTGAGGGGGAAATAA-3’, HACNS50 R 5’-TTATTTCCTTCAGGCCCTTG-3’, and cloned into an Hsp68-lacZ reporter vector as previously described^95^. Generation of transgenic mice at the Yale University Transgenic Mouse Facility and embryo staining were carried out as previously described^95^. We required reporter gene expression in a given structure to be present in at least three independent transgenic embryos as assessed by two researchers to be considered reproducible.

### Circularized Chromosome Conformation Capture with Sequencing (4C-Seq)

All animal work was done in accordance with approved University of Connecticut Health Center IACUC protocols. 4C-seq was performed according to van de Werken et al. (2012)^76^ with modifications for tissue. Input mouse embryonic craniofacial and brain tissue from the same litter was fixed and nuclei isolated following homogenization with a dounce tissue grinder as described^79^. Each replicate consists of tissue from an individual litter. Subsequent digestion and ligation steps were followed from van de Werken et al. (2012)^76^. Chromatin was digested sequentially with NlaIII and DpnII. Amplification of final libraries was performed with primers selected using a primer database generated for NlaIII/DpnII digestion as previously described^76^. The sequences added to these primers were modified to allow hybridization to NextSeq 500 flow cells and split across two sets of primers to improve efficiency and allow for dense multiplexing (Table S9).

### 4C-seq Data Analysis

4C-seq libraries were sequenced for 75 cycles using the NextSeq500 (Illumina). Fastq files were demultiplexed by barcode yielding Fastq files for each tissue replicate. Tissue replicate Fastq files were further demultiplexed by viewpoint using Cutadapt (v1.8.3)^96^. Trimmed reads were uniquely aligned to mm9 using bowtie2^82^. Significant interactions in craniofacial tissue were assessed using r3Cseq^97^ with a modification allowing a larger viewing window near the viewpoint (https://github.com/cotneylab/r3Cseq) and using brain as a control. The significant interactions are represented in the accompanying track hub as bigBed files. The location of the viewpoint and sequenced interacting fragment are denoted with thick bars. A thin bar is included to denote the connection between the viewpoint and the distal sites.

## Supplemental Figure and Table Legends

**Supplemental figures and tables can be obtained from FigShare:**

**10.6084/m9.figshare.4954202**

**Supplementary Figure 1. Detailed Histone Modification Profiles in Human Craniofacial Development. a.** Heatmap and hierarchical clustering of pairwise Pearson correlations for 114 individual histone modification profiles from human craniofacial tissues. Darker orange indicates positive correlation between datasets. Enlarged from **Fig. 2a** to include sample details, showing samples cluster closely by histone mark. **b.** Correlation of only H3K27ac data contained in the area boxed in black in part **a**. Heatmap and hierarchical clustering show that the samples cluster well into groups by early or late stage of development.

**Supplementary Figure 2. Complete Histone Modification Profiles in Human Craniofacial Development** Genomic feature annotations identified by peak calls from six histone modification profiles from all craniofacial samples, across all Carnegie stages, plotted as cumulative percentage of total peaks. Peak enrichments and genomic annotations were performed using HOMER^83^.

**Supplementary Figure 3. Imputed Histone Modification Profiles in Human Craniofacial Development. a.** Heatmap and hierarchical clustering of pairwise Pearson correlations for imputed histone modification profiles from human craniofacial tissues. Darker orange indicates positive correlation between datasets. **b.** Heatmap and hierarchical clustering of pairwise Pearson correlations for imputed and primary histone modification profiles from human craniofacial tissues. Darker orange indicates positive correlation between datasets.

**Supplementary Figure 4. Imputation of Craniofacial Epigenomic Signals and Chromatin State Segmentation in the 15-State (Primary) and 18-State (Auxiliary) ChromHMM models. a.** Numbers of individual chromatin state segments identified by each of the color-coded 15 states of chromatin activity based on imputed epigenomic signals for each of the 21 tissue samples profiled. **b**. Comparison of cumulative percentage of each chromatin state between craniofacial samples profiled here and 127 segmentations generated by Roadmap Epigenome^22^. **c**. Mean numbers of segments annotated in each of the 15 states across 21 craniofacial samples (orange) and 127 Roadmap Epigenomes (gray). **d.** Mean percentages of segments annotated in each of the 15 states across 21 craniofacial samples (orange) and 127 Roadmap Epigenomes (gray). **e.** Same as in panel **a**, but for 18-State Model. **f.** Same as in panel **b**, but for 18-State Model. **g.** Same as in panel **c**, but for 18-State Model. **h.** Same as in panel **d**, but for 18-State Model. Error bars represent standard deviation. Overall chromatin state segmentation in craniofacial samples identifies similar numbers and percentages of each of the states published by Roadmap Epigenome^22^.

**Supplementary Figure 5. All Enhancers Tested for Craniofacial Activity** All enhancers identified and tested by this study from the Vista Enhancer Browser. Enhancers with hs prefix indicated the human genomic sequence was tested while those with the mm prefix indicate that the orthologous sequence from mouse identified by this study was tested.

**Supplementary Figure 6. H3K27ac Signal at Enhancer Segments Allows for Correlation by Tissue Type. a.** Heatmap and hierarchical clustering of pairwise comparisons of H3K27ac signals at all enhancer segments in our craniofacial data and the 127 samples from Roadmap Epigenome. Red coloring indicates positive correlation between datasets, blue indicates less correlation. **b.** Principal component analyses of the first four component dimensions of H3K27ac signals in a serial progressive fashion (i.e PC1 vs PC2, PC2 vs PC3, etc.). Samples are color coded by tissue type.

**Supplementary Figure 7. Identification of Craniofacial-specific Enhancers Flanking *MSX2***. Enhancer states annotated by the 25-state model that are found only in craniofacial tissue but not the 127 samples from Roadmap Epigenome are located upstream and downstream of *MSX2*, a gene implicated in multiple craniofacial abnormalities. The enhancer states fall within a region of conservation and are supported at top by ChIP signals from a single human craniofacial tissue sample.

**Supplementary Figure 8. Integration of CL/P GWAS Data Places SNPs within Craniofacial-specific Enhancers. a.** Enrichment analysis identified orofacial cleft GWAS tag SNPs preferentially among craniofacial tissue. **b.** Enhancer state analysis permits placement of a potentially causative allele for non-syndromic CL/P (rs642961) within a predicted early development enhancer state. This enhancer state is located between *IRF6* and *DIEXF* and may influence expression of *IRF6*. **c.** The orofacial cleft-associated tag SNP rs745080 resides within the intron of *TXNDC16*, which is marked by a craniofacial-specific enhancer state.

**Supplementary Figure 9. Clustering Identifies Similar Functional Enrichment to Self-Organizing Maps a.** K-means clustering performed directly on the matrix of H3K27ac signals, using the same number of clusters utilized for the self-organizing map (199), showed distinct H3K27ac activation in craniofacial tissues (enlarged section on the right). **b.** Analysis of the sequence content of enhancer clusters most specific for craniofacial activity identified enrichments of motifs for the *HOX*, *LHX*, *MSX* and *DLX* families of transcription factors.

**Supplementary Figure 10. Incorporation of Topologically Associated Domain Structure and Clinical Case Suggests Interaction Between Distal Enhancer and HoxA Region.** Hi-C data from HUVEC visualized using the Hi-C browser (http://promoter.bx.psu.edu/hi-c/view.php) indicates an interaction between the *HOXA* cluster and an intergenic region approximately 1Mb away from the anterior side. A deletion covering this region and notably leaving the *HOXA* cluster intact has been described in a patient with facial dysgenesis^78^ (indicated in purple). The intergenic region contains a 450kb enhancer state, as represented by the 15-, 18- and 25-state ChromHMM model of craniofacial data, with regions of high conservation. Four viewpoints used in 4C-seq are indicated, flanking the region of interest. Seven enhancers with craniofacial activity are located within this region, indicated by black bars and representative images. Enhancers mm403-407 and hs1600 were tested by the Vista Enhancer Browser^50^, HACNS50 was tested independently.

**Supplementary Figure 11. Distal Regulatory Region Shares Chromatin State in Mouse.** ChIP-Seq data from Mouse Encode for embryonic day 11.5 facial prominence display a conserved set of chromatin marks in the region distal to the *HOXA* cluster suggesting a conserved function in mouse craniofacial development.

**Supplementary Figure 12. Syntenic Block Near *HOXA* Cluster.** A comparison of a 10 Mb window around the *HOXA* cluster on human Chromosome 7 shows synteny with mouse Chromosome 6. In addition to the preservation of gene order, there is also preservation of a large non-coding region distal to the anterior side of the *HOXA* cluster in mouse.

**Supplementary Figure 13. Targeted Sequencing of 13 Loci Identified by GWAS Studies to be Important In Craniofacial Development Misses a Regulatory Region in *BMP4***. The study by Leslie et al.^77^ performed targeted sequencing of a region of ∼60 kb surrounding the BMP4 gene (black bar at top of figure). This region excluded a region immediately adjacent (outlined by green box) identified as an enhancer by the 25-State, Imputed ChromHMM model in all 21 craniofacial tissues analyzed.

**Supplementary Table 1.** Table showing, for each sample, for each mark, the number of total sequencing reads, the number of uniquely mapped reads, the number of multi-mapped reads, percentage of mapped reads, and percentage of uniquely mapped reads. Total numbers, in billions, and means, in millions, are displayed in bottom two rows of the table.

**Supplementary Table 2.** Table showing all enhancer segments in craniofacial epigenomic atlas that were never annotated as any type of enhancer state in all of Roadmap Epigenome.

**Supplementary Table 3.** Motifs identified for enrichment of transcription factor binding sites for novel craniofacial-specific enhancers found in this study. Consensus motif sequence, p-values, q-values, number of target sequences with the motif, percent of target sequences with the motif, number of background sequences with the motif, and percent of all background sequences with the motif are all indicated.

**Supplementary Table 4.** Functional categories with significant enrichment based on assignment of craniofacial-specific enhancers to the nearest gene, using Genomic Regions Enrichment of Annotations Tool (GREAT)^58^.

**Supplementary Table 5.** Regions containing tag SNPS identified in craniofacial tissue and associated with orofacial clefting, including gene assignments where applicable. Of note is a tag SNP in the noncoding region between *IRF6* and *DEXIF*, as well as an intronic sequence within the *TXNDC16* gene.

**Supplementary Table 6.** Functional categories with significant enrichment in the clusters of craniofacial-specific enhancers (annotated to the the nearest gene) obtained from K-means clustering on the matrix of H3K27ac signals using the same number of clusters utilized for the self-organizing map, using Genomic Regions Enrichment of Annotations Tool (GREAT)^58^.

**Supplementary Table 7.** Sheet1: 582 regions identified across the genome as containing significant fractions of bases annotated as craniofacial enhancers; start position, end position, and fraction of bases annotated as an enhancer state are shown. The second sheet shows individual window analysis with fold enrichment versus randomized enhancer segmentations and permutation p-values.

**Supplementary Table 8.** Functional categories with significant enrichment based on assignment of enriched enhancer windows to the nearest gene, using Genomic Regions Enrichment of Annotations Tool (GREAT)^58^.

**Supplementary Table 9.** Primers used for 4C-Seq analysis in mouse craniofacial tissue.

